# Input-specific control of interneuron numbers in nascent striatal networks

**DOI:** 10.1101/2021.12.01.470575

**Authors:** Varun Sreenivasan, Eleni Serafeimidou-Pouliou, David Exposito-Alonso, Kinga Bercsenyi, Clémence Bernard, Sung-Eun Bae, Fazal Oozeer, Alicia Hanusz-Godoy, Robert Edwards, Oscar Marín

**Affiliations:** Centre for Developmental Neurobiology, Institute of Psychiatry, Psychology and Neuroscience, King’s College London, London SE1 1UL, United Kingdom; MRC Centre for Neurodevelopmental Disorders, King’s College London, London SE1 1UL, United Kingdom; Department of Physiology and Department of Neurology, School of Medicine, University of California San Francisco, United States of America

## Abstract

The assembly of functional neuronal circuits requires appropriate numbers of distinct classes of neurons, but the mechanisms through which their relative proportions are established remain poorly defined. Investigating the mouse striatum, here we found that the two most prominent subtypes of striatal interneurons, parvalbumin-expressing (PV+) GABAergic and cholinergic (ChAT+) interneurons, undergo extensive programmed cell death between the first and second postnatal weeks. Remarkably, the survival of PV+ and ChAT+ interneurons is regulated by distinct mechanisms mediated by their specific afferent connectivity. While long-range cortical inputs control PV+ interneuron survival, ChAT+ interneuron survival is regulated by local input from the medium spiny neurons. Our results identify input-specific circuit mechanisms that operate during the period of programmed cell death to establish the final number of interneurons in nascent striatal networks.

---

There are hundreds of different types of projection neurons and interneurons in the mammalian nervous system, but the various cellular components of each brain structure arise during development in very precise proportions. This is largely achieved by an evolutionarily conserved strategy based on the initial overproduction of cells and the subsequent elimination of a fraction of them through the process of programmed cell death (Oppenheim, 1991; Raff et al., 1993).

In the neocortex, the relative proportion of excitatory pyramidal neurons and inhibitory GABAergic interneurons is similar across cortical areas and even species (DeFelipe et al., 2002; Hendry et al., 1987). In mice, pyramidal cells and interneurons undergo extensive programmed cell death during early postnatal development to adjust their final ratio (Wong and Marín, 2019). Cortical interneurons appear to be intrinsically programmed to undergo cell death unless they sustain a certain level of activity during early postnatal development (Denaxa et al., 2018; Priya et al., 2018; Southwell et al., 2012; Wong et al., 2018). This mechanism guarantees that only interneurons that receive sufficient input from pyramidal cells are retained (Duan et al., 2020; Wong et al., 2018).

The striatum is another brain structure that contains both projection neurons and interneurons. However, it is unique in its almost complete lack of glutamatergic neurons. The principal neurons of the striatum, called medium spiny neurons (MSNs), are GABAergic and constitute 95% of all striatal neurons (Graveland and DiFiglia, 1985). Single-cell transcriptomic analyses have identified several populations of striatal interneurons in the mouse (Muñoz-Manchado et al., 2018), including two major types for which there is abundant functional, morphological, and electrophysiological information: (1) parvalbumin-expressing (PV+) GABAergic interneurons; and (2) choline acetyltransferase expressing (ChAT+) cholinergic interneurons (Kawaguchi, 1993). Like many cortical interneurons, striatal PV+ and ChAT+ interneurons derive from progenitor cells in the medial ganglionic eminence (MGE) and preoptic area (POA) of the embryonic telencephalon (Marín et al., 2000), but it is presently unknown whether these cells also undergo programmed cell death during early postnatal development. In addition, the absence of local excitatory neurons raises the question of how the appropriate ratio of projection neurons and interneurons is established in the striatum and whether the cellular mechanisms underlying this process are universal across the brain. Here we found that striatal interneurons undergo extensive programmed cell death, a process that is specifically regulated by their afferent connectivity during a critical window of early postnatal development. Our experiments reveal that activity-dependent, input-specific control of programmed cell death regulates interneuron numbers in the striatum.

## Results

### Striatal interneurons undergo programmed cell death

To investigate whether striatal interneurons undergo programmed cell death during postnatal development, we generated *Nkx2-1-Cre;RCL^tdTomato^* mice in which MGE/POA-derived interneurons are irreversibly labeled from early embryonic development and quantified the number of tdTomato-expressing interneurons in the striatum using stereological methods. Between postnatal days (P) 2 and P 21 (Fig. 1A) we observed a dramatic decrease in the total number of striatal interneurons. By the end of the third postnatal week, the total number of MGE/POA-derived interneurons in the striatum reduced to nearly half the number found around birth, with the most significant variation occurring between P5 and P10 (Fig. 1B and C).

**Fig. 1.**
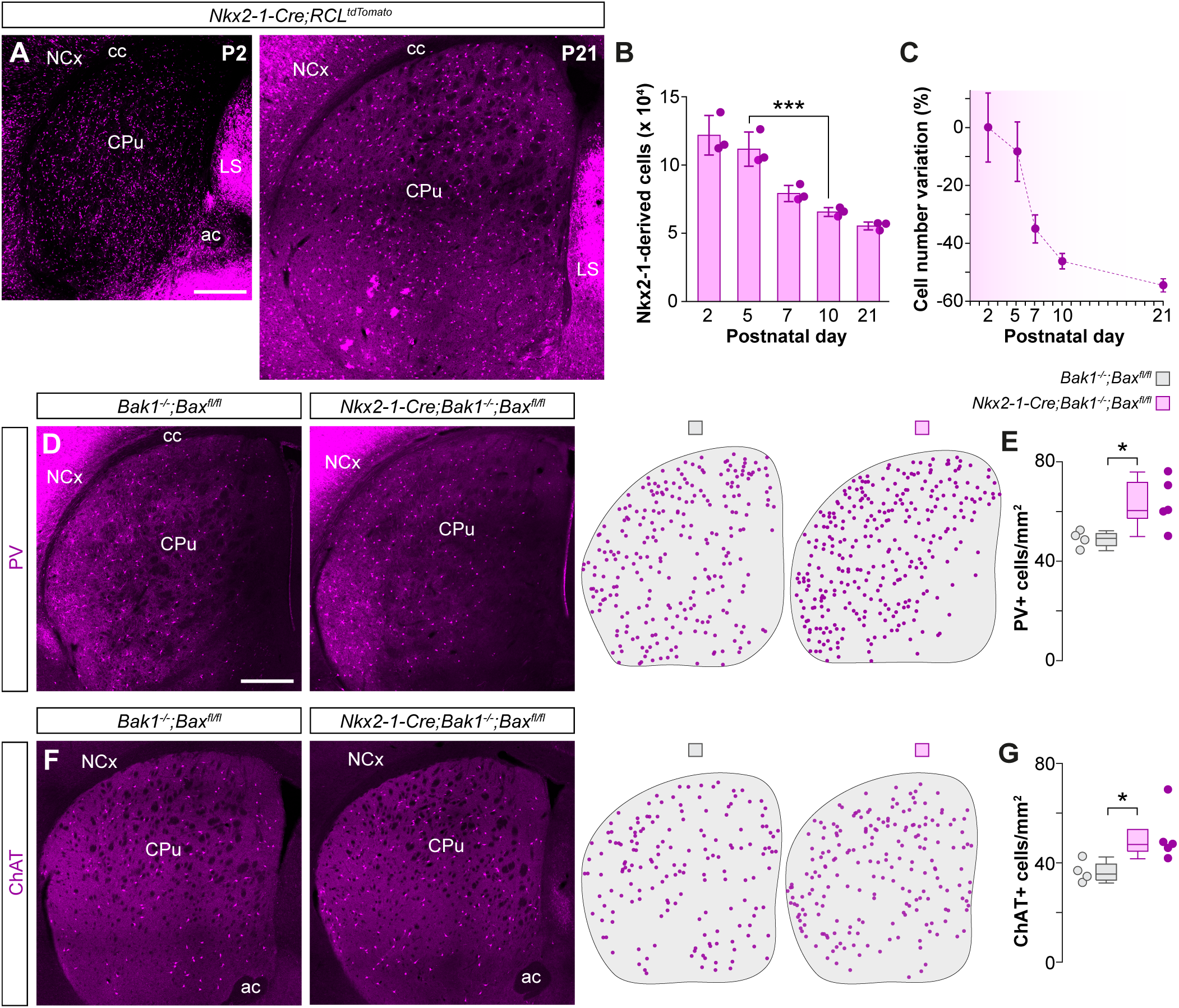
Striatal interneurons undergo programmed cell death during early postnatal development. **(A)** Coronal sections through the striatum of *Nkx2-1-Cre;RCL^TdTomato^* mice at P2 and P21. **(B)** Stereological estimates of the total number of tdTomato+ neurons in the striatum of *Nkx2-1-Cre;RCL^TdTomato^* mice at different postnatal stages (*n* = 3 mice per postnatal stage); one-way ANOVA, \*\*\**p* < 0.001; P2 vs. P5: Tukey-Kramer HSD test, *p* > 0.05; P5 vs. P10: Tukey-Kramer HSD test, \*\*\**p* < 0.001. **(C)** Percentage variation in the number of tdTomato+ cells at different postnatal stages relative to the average number of tdTomato+ cells at P2 (right). **(D and F)** Coronal sections through the striatum of control *Bak1^-/-^;Bax^fl/fl^* and mutant *Nkx2-1-Cre;Bak1^-/-^;Bax^fl/fl^* mice immunostained for PV (D) and ChAT (F). The schematic dot maps indicate the locations of striatal interneurons in each case. **(E and G)** Quantification of PV+ (E) and ChAT+ (G) interneuron density in the striatum of control *Bak1^-/-^;Bax^fl/fl^* (*n* = 4) and mutant *Nkx2-1-Cre;Bak1^-/-^;Bax^fl/fl^* (*n* = 5) mice. PV+ interneurons: two-tailed unpaired Student’s *t*-test, \**p* < 0.05. ChAT+ interneurons: Wilcoxon’s rank-sum test, \**p* < 0.05. Data in panel B are shown as bar plots and data in panels E and G as box plots. Adjacent data points indicate the stereological estimate in each mouse (B) or the average cell density in each animal (E and G). Error bars indicate standard deviation. Scale bars, 500 *μ*m.

We next sought to determine whether different populations of striatal MGE/POA-derived interneurons undergo programmed cell death at comparable rates. We hypothesized that like their cortical counterparts (Southwell et al., 2012; Wong et al., 2018), striatal interneurons might require the concerted action of the BCL2 family genes *Bax* and *Bak1* to undergo cell death. Consequently, removing *Bax* and *Bak1* in striatal interneurons would reveal how this process specifically affects distinct interneuron classes. To test this hypothesis, we generated *Nkx2-1-Cre;Bak1^-/-^;Bax^fl/fl^* mutant mice and quantified the density of PV+ and ChAT+ striatal interneurons. Suppressing cell death in MGE/POA-derived cells led to a ~30% increase in the density of PV+ interneurons (Fig. 1D and E) and a ~40% increase in the density of ChAT+ interneurons (Fig. 1F and G), indicating that both PV+ and ChAT+ interneuron numbers are strongly modulated during early postnatal development.

### Cortical pyramidal neurons regulate striatal interneuron survival

The survival of cortical GABAergic interneurons is regulated by the number and activity of local excitatory neurons (Wong et al., 2018). The striatum lacks an equivalent population of local excitatory neurons, but its main source of excitatory input originates from the neocortex along with prominent input from the thalamus (Johansson and Silberberg, 2020; Klug et al., 2018; Wall et al., 2013). To test if long-range cortical inputs regulate striatal interneuron survival, we first asked whether preventing cortical excitatory neurons from undergoing programmed cell death – thereby artificially increasing their number – would, in turn, change the survival of striatal interneurons. To this end, we quantified the density of PV+ and ChAT+ interneurons in *Nex^Cre/+^;Bak1^-/-^;Bax^fl/fl^* mice, in which the programmed cell death of cortical pyramidal cells is specifically abolished (Wong et al., 2018). Artificially increasing the number of pyramidal neurons during development led to a ~20% increase in the density of PV+ interneurons throughout the striatum (Fig. 2A and C). In contrast, we observed no changes in the density of ChAT+ interneurons (Fig. 2B and D).

**Fig. 2.**
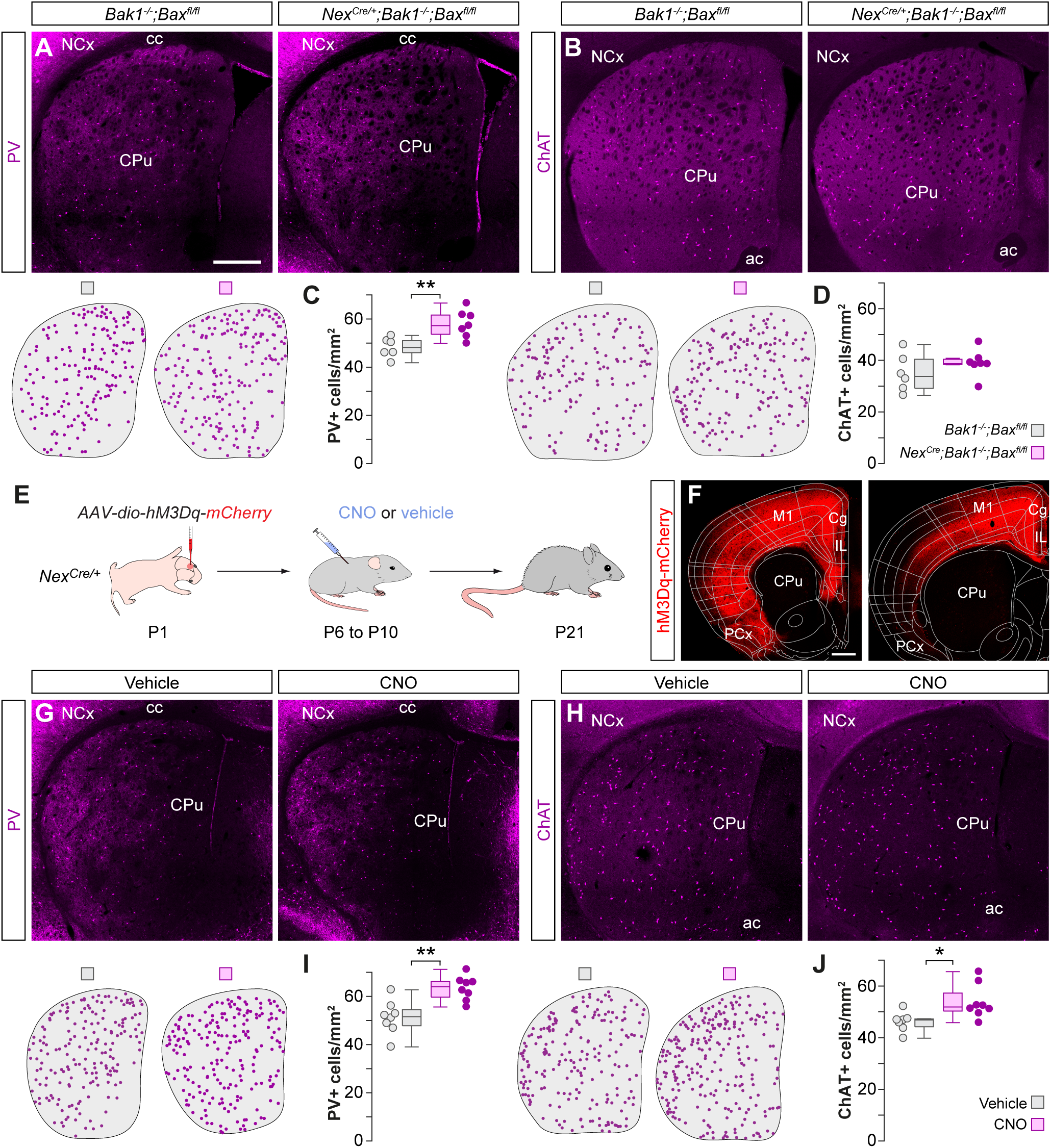
Cortical gain of function rescues striatal PV+ and ChAT+ interneurons. **(A and B)** Coronal sections through the striatum of control *Bak1^-/-^;Bax^fl/fl^* mice and mutant *Nex^Cre/+^;Bak1^-/-^;Bax^fl/fl^* mice immunostained for PV (A) and ChAT (B). The schematic dot maps indicate the locations of striatal interneurons in each case. **(C and D)** Quantification of PV+ (C) and ChAT+ (D) interneuron density in the striatum of control *Bak1^-/-^;Bax^fl/fl^* (*n* = 6) and mutant *Nex^Cre/+^;Bak1^-/-^;Bax^fl/fl^* (*n* = 7) mice. PV+ interneurons: two-tailed unpaired Student’s *t*-test, \*\**p* < 0.01. ChAT+ interneurons: two-tailed unpaired Student’s *t*-test, *p* = 0.27. **(E)** Schematic of experimental design. **(F)** Coronal sections through the brain at ~1.5 mm (left) and ~0.5 mm (right) anterior to Bregma showing the extent of the viral infection. The images have been overlaid with the corresponding coronal maps. **(G and H)** Coronal sections through the striatum of vehicle and CNO-treated *Nex^Cre/+^* mice immunostained for PV (G) and ChAT (H). The schematic dot maps indicate the locations of striatal interneurons in each case. **(I and J)** Quantification of PV+ (I) and ChAT+ (J) interneuron density in the striatum of vehicle (*n* = 8 for PV and 7 for ChAT) and CNO-treated (*n* = 8) *Nex^Cre/+^* mice. PV+ interneurons: two-tailed unpaired Student’s *t*-test, \*\**p* < 0.01. ChAT+ interneurons: twotailed unpaired Student’s *t*-test, \**p* < 0.05. Data in panels C, D, I and J are shown as box plots and the adjacent data points indicate the average cell density in each animal. Scale bars, 500 *μ*m.

The previous results revealed that the survival of striatal PV+ interneurons depends on the number of cortical pyramidal cells. PV+ interneurons preferentially receive synaptic input from cortico-striatal projections (Johansson and Silberberg, 2020), which might explain the differences observed among interneuron subtypes. Therefore, we investigated whether increasing the activity of cortical pyramidal cells during the period of striatal interneuron cell death would affect their survival. To this end, we transiently increased the activity of pyramidal cells using the chemogenetic actuator hM3Dq (Roth, 2016). We expressed an adeno-associated virus (AAV) encoding Cre dependent hM3Dq in the frontal cortex of *Nex^Cre/+^* mice at P1 (Fig. 2E and F). We then injected pups with the ligand clozapine –N-oxide (CNO) or vehicle twice daily between P6 and P10 and examined the distribution of striatal interneurons at P21. Enhancing pyramidal cell activity during the period of striatal interneuron cell death led to a robust ~24% increase in the density of PV+ interneurons (Fig. 2G and I) and a more modest increase of ~17% in the density of ChAT+ interneurons (Fig. 2H and J).

To rule out the possibility that the observed increase in PV+ interneuron density is due to changes in PV levels and not in actual interneuron numbers, we quantified the density of striatal Nkx2-1+ cells that do not express ChAT. At P21, striatal Nkx2-1+/ChAT-cells comprise all PV+ interneurons and a very small percentage of somatostatin neurons (Marín et al., 2000). We found an increase in Nkx2-1+/ ChAT-interneuron density in CNO-treated mice compared to controls (Fig. S1B-C) indicating that the increased density of striatal PV+ interneurons is likely due to increased survival and not to activity-dependent changes in PV levels.

### Paradoxical effects of cortical dysfunction on striatal interneuron survival

We next investigated whether the modulation of striatal interneuron numbers by cortical pyramidal neurons is bi-directional. Based on our previous results, we hypothesized that decreasing the number or activity of pyramidal neurons, during the period of programmed cell death, would negatively impact PV+ and ChAT+ interneuron survival. To test this hypothesis, we first generated *Rbp4-Cre;Stxbp1^fl/fl^* mice, in which syntaxin-binding protein 1 (also known as Munc18-1) is removed from Rbp4-expressing (Rbp4+) layer 5 pyramidal neurons in the neocortex. Deleting *Stxbp1* blocks neurotransmission, which is followed by rapid apoptosis of the cell (Verhage et al., 2000). By P6, we found that mutant mice contained significantly fewer Ctip2+ layer 5 pyramidal cells than control mice in primary somatosensory barrel cortex and primary motor cortex (Fig. S2A-B). We then assessed whether the loss of cortico-striatal projection neurons impacted the survival of striatal interneurons. Consistent with our hypothesis, we found that mutant mice contained ~12% fewer striatal PV+ interneurons than controls at P21 (Fig. 3A and C). Unexpectedly, we also observed a ~20% increase in the density of striatal ChAT+ interneurons in mutants compared to control mice (Fig. 3B and D).

**Fig. 3.**
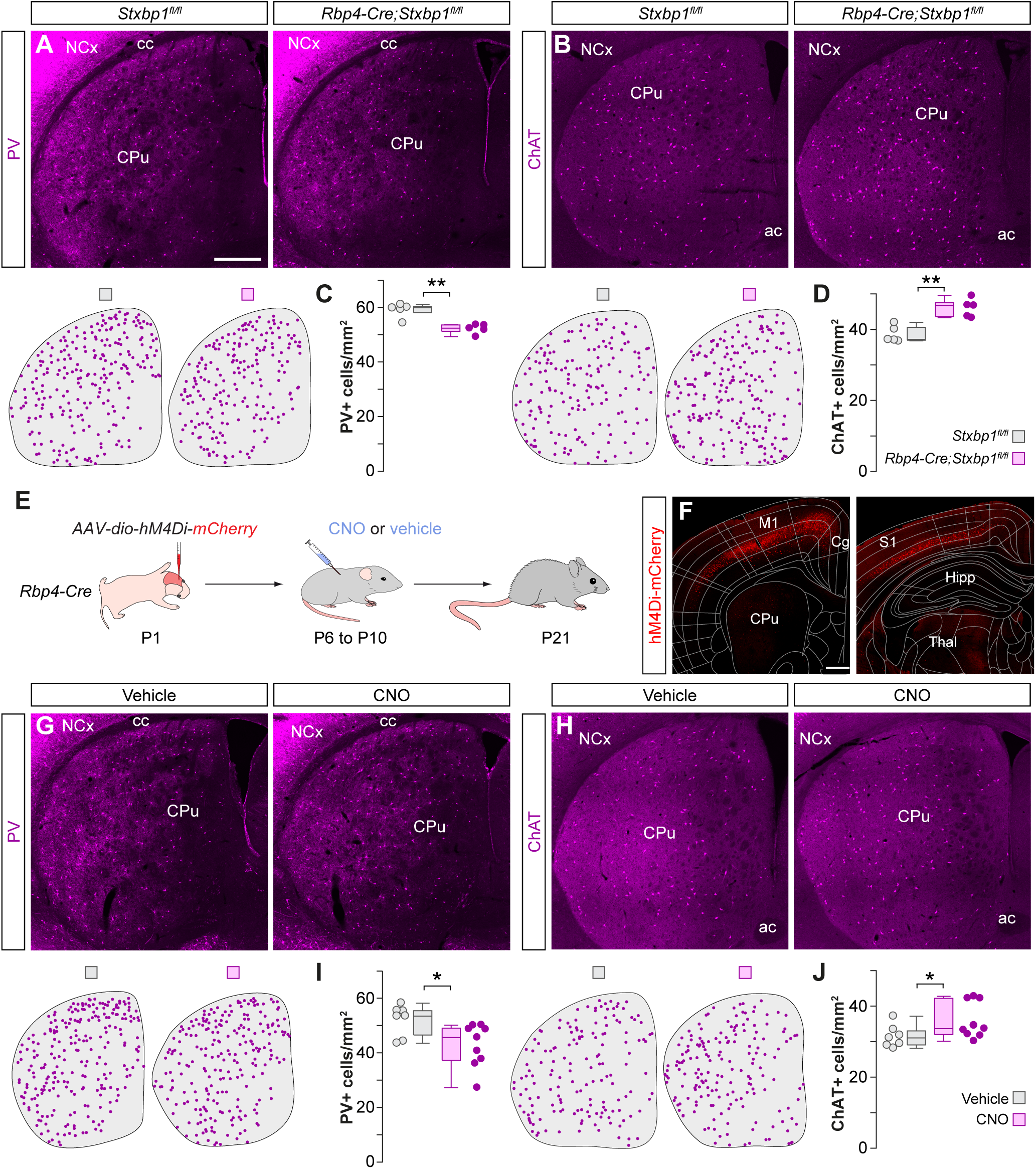
Cortical dysfunction differentially impacts the survival of striatal PV+ and ChAT+ interneurons. **(A and B)** Coronal sections through the striatum of control *Stxbp1^fl/fl^* mice and mutant *Rbp4-Cre;Stxbp1^fl/fl^* mice immunostained for PV (A) and ChAT (B). The schematic dot maps indicate the locations of striatal interneurons in each case. **(C and D)** Quantification of PV+ (C) and ChAT+ (D) interneuron density in the striatum of control *Stxbp1^fl/fl^* (*n* = 5) and mutant *Rbp4-Cre;Stxbp1^fl/fl^* (*n* = 5) mice. PV+ interneurons: Wilcoxon’s rank-sum test, \*\**p* < 0.01. ChAT+ interneurons: two-tailed unpaired Student’s *t*-test, \*\**p* < 0.01. **(E)** Schematic of experimental design. **(F)** Coronal sections through the brain at ~1 mm anterior (left) and ~2 mm posterior (right) to Bregma showing the extent of the viral infection. The images have been overlaid with the corresponding coronal maps. **(G and H)** Coronal sections through the striatum of vehicle- and CNO-treated *Rbp4-Cre* mice immunostained for PV (G) and ChAT (H). The schematic dot maps indicate the locations of striatal interneurons in each case. **(I and J)** Quantification of PV+ (I) and ChAT+ (J) interneuron density in the striatum of vehicle (*n* = 7) and CNO-treated (*n* = 9) *Rbp4-Cre* mice. PV+ interneurons: twotailed unpaired Student’s *t*-test, \**p* < 0.05. ChAT+ interneurons: Wilcoxon’s rank-sum test, \**p* = 0.05. Data in panels C, D, I and J are shown as box plots and the adjacent data points indicate the average cell density in each animal. Scale bars, 500 *μ*m.

Next, we tested whether decreasing the activity of pyramidal neurons has similar effects on striatal interneuron survival as reducing their number. We expressed Cre-dependent AAVs encoding the chemogenetic actuator hM4Di (Roth, 2016) throughout the neocortex of *Rbp4-Cre* mice at P1 (Fig. 3E and F). We then injected pups with CNO or vehicle thrice daily between P6 and P10 and examined the distribution of striatal interneurons at P21. We found a significant decrease of ~18% in the density of PV+ interneurons in CNO-treated mice compared to controls (Fig. 3G and I). Interestingly, we again observed a significant increase of ~14% in the density of striatal ChAT+ interneurons (Fig. 3H and J). Altogether, these experiments demonstrated that the survival of striatal PV+ interneurons can be bidirectionally modulated by the number and activity of cortical pyramidal neurons. These experiments also revealed paradoxical effects on ChAT+ interneurons since both bidirectional manipulations led to an increase in the survival of this population.

### PV+ interneuron survival requires glutamatergic neurotransmission

In the neocortex, the activity of individual MGE interneurons correlates with their survival (Wong et al., 2018) and lowering their activity in a cell-autonomous manner reduces their ability to survive (Duan et al., 2020). Since the activity of any given neuron is largely derived from glutamatergic inputs, we first tested whether striatal interneuron survival depends on glutamatergic neurotransmission from the cortex. To delete the vesicular glutamate transporters Vglut1 and Vglut2, we crossed double heterozygous *Vglut1* mutant and floxed *Vglut2* mice, and injected Cre-expressing AAVs in the neocortex of the resulting control (*Vglut1^+/+^;Vglut2^+/+^*) and double mutant (*Vglut1^-/-^;Vglut2^fl/fl^*) animals at P1 (Fig. 4A and B). We found a very strong decrease of ~70% in the density of PV+ interneurons in Cre-injected *Vglut1^-/-^;Vglut2^fl/fl^* mice compared to controls in the striatum (Fig. 4C and E) as well as in the cortex (Fig. S3A-B) at P21. In contrast, the loss of cortical glutamate led to a significant increase of ~21% in the survival of striatal ChAT+ interneurons (Fig. 4D and F). These results revealed that glutamate neurotransmission in cortico-striatal neurons is crucial for the survival of striatal PV+ interneurons but not ChAT+ interneurons.

**Fig. 4.**
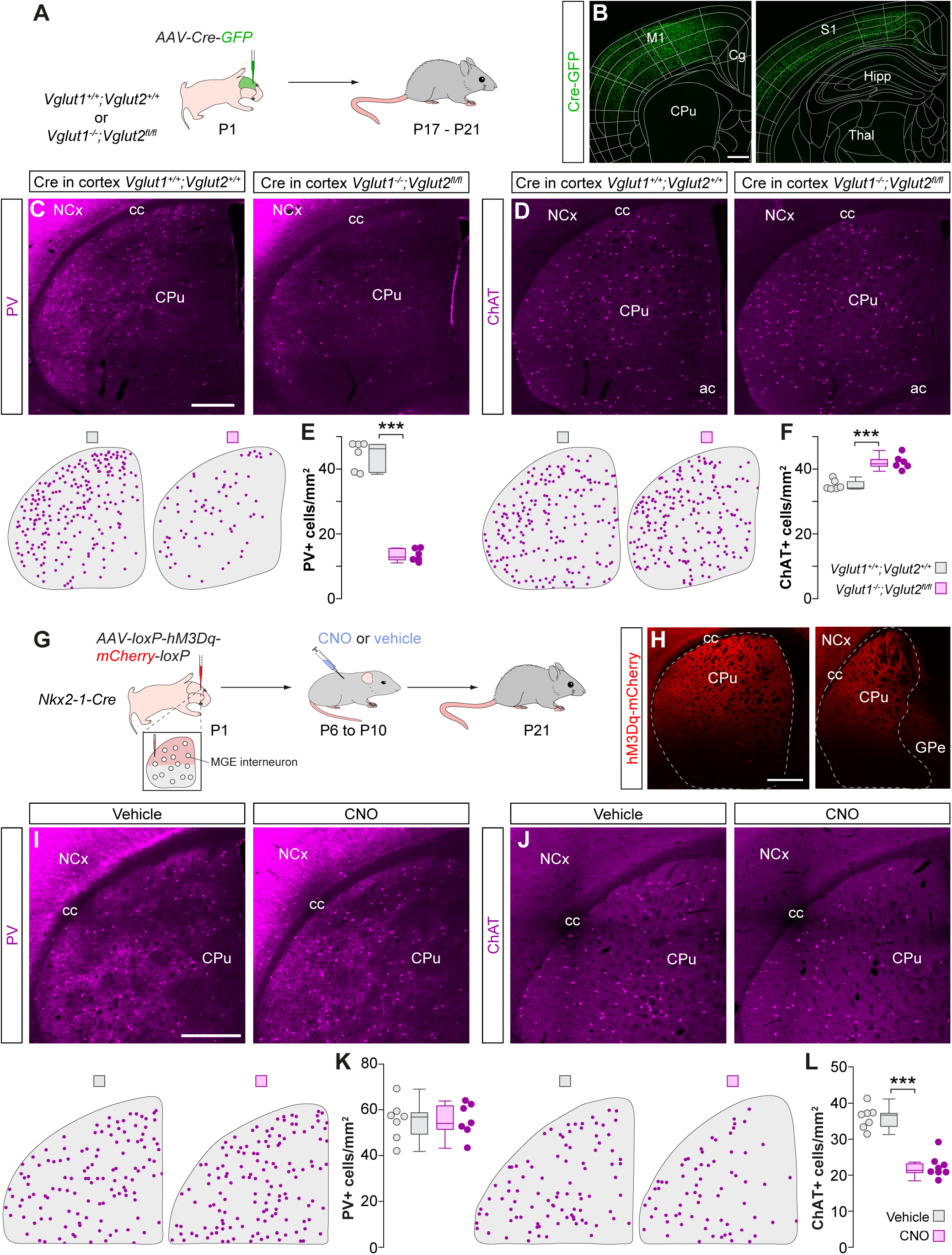
Input specific mechanisms for striatal interneuron survival. **(A)** Schematic of experimental design. **(B)** Coronal sections through the brain at ~1 mm anterior (left) and ~2 mm posterior (right) to Bregma showing the extent of the viral infection. The images have been overlaid with the corresponding coronal maps. **(C and D)** Coronal sections through the striatum of Cre injected control *Vglut1^+/+^;Vglut2^+/+^* and double-mutant *Vglut1^-/-^;Vglut2 ^fl/fl^* mice immunostained for PV (C) and ChAT (D). The schematic dot maps indicate the locations of striatal interneurons in each case. **(E and F)** Quantification of PV+ (E) and ChAT+ (F) interneuron density in the striatum of Cre injected control *Vglut1^+/+^;Vglut2^+/+^* (*n* = 6) and double-mutant *Vglut1^-/-^;Vglut2 ^fl/fl^* (*n* = 6) mice. PV+ interneurons: two-tailed unpaired Student’s *t*-test, \*\*\**p* < 0.001. ChAT+ interneurons: two-tailed unpaired Student’s *t*-test, \*\*\**p* < 0.001. **(G)** Schematic of experimental design. **(H)** Coronal sections through the striatum showing the extent of the viral infection. Dashed lines indicate the boundary of the striatum. **(I and J)** Coronal sections through the dorsal striatum of vehicle- and CNO-treated *Nkx2-1-Cre* mice immunostained for PV (I) and ChAT (J). The schematic dot maps indicate the locations of striatal interneurons in each case. **(K and L)** Quantification of PV+ (K) and ChAT+ (L) interneuron density in the dorsal striatum of vehicle (*n* = 7) and CNO-treated (*n* = 7 for PV and 8 for ChAT) *Nkx2-1-Cre* mice. PV+ interneurons: two-tailed unpaired Student’s *t*-test, *p* = 0.9. ChAT+ interneurons: two-tailed unpaired Student’s *t*-test, \*\*\**p* < 0.001. Data in panels E, F, K and L are shown as box plots and the adjacent data points indicate the average cell density in each animal. Scale bars, 500 *μ*m.

### ChAT+ interneuron survival depends on MSN activity

Striatal ChAT+ interneurons receive relatively few direct inputs from pyramidal cells (Johansson and Silberberg, 2020; Lapper and Bolam, 1992), which suggests that the paradoxical effects on ChAT+ interneuron survival, caused by manipulating the cortex, might be indirect. We hypothesized that the survival of ChAT+ interneurons might be controlled by the activity of the MSNs since ChAT+ interneurons receive particularly strong inputs from these cells (Chuhma et al., 2011). We tested this hypothesis by injecting AAVs expressing hM3Dq in the striatum of *Nkx2-1-Cre* mice at P1 (Fig. 4G and H). To prevent the expression of hM3Dq in PV+ and ChAT+ interneurons, we flanked the expression construct with loxP sequences to excise it out in Cre expressing populations. We then injected pups with CNO or vehicle thrice daily between P6 and P10 and examined the distribution of striatal interneurons at P21. Activating MSNs during the period of striatal interneuron cell death did not impact the survival of PV+ interneurons (Fig. 4I and K). In contrast, we observed a prominent ~38% decrease in the density of ChAT+ interneurons (Fig. 4J and L) suggesting that MSN activity, during this period, negatively impacts the survival of ChAT+ interneurons.

## Discussion

Our study demonstrates that the neocortex can remotely influence the establishment of neural circuits in another region of the brain, the striatum. We found that the two most prominent types of striatal interneurons, PV+ GABAergic neurons and ChAT+ cholinergic neurons, undergo substantial programmed cell death during a short period of early postnatal development in the mouse. Their survival is under the control of specific afferent inputs during the cell death period. The final number of PV+ interneurons is established by long-range cortical inputs, while local inputs from medium spiny neurons regulate the final density of ChAT+ interneurons. Our results reveal circuit-based rules for establishing interneuron ratios and point to activity-dependent, input-specific mechanisms as the main determinant of the final number of interneurons in nascent striatal networks (Fig. S4).

### Long-range control of striatal PV+ interneuron survival

Striatal PV+ interneurons mediate inhibition onto neighboring MSNs (Gittis et al., 2010; Koos and Tepper, 1999; Straub et al., 2016) and they are primarily driven by long-range excitatory inputs from the cortex and the thalamus (Johansson and Silberberg, 2020; Ramanathan et al., 2002; Rudkin and Sadikot, 1999). We found that modifying the number or activity of cortical pyramidal neurons bi-directionally regulates the survival of these interneurons. Given the key role these interneurons play in controlling MSN activity in response to cortical input (Gittis et al., 2010; Koos and Tepper, 1999; Mallet et al., 2005; Straub et al., 2016), it seems logical that pyramidal cells are directly involved in establishing their final numbers.

The neocortex controls the survival of striatal PV+ interneurons through glutamatergic neurotransmission. Preventing exocytotic glutamate release from cortical pyramidal cells led to the elimination of nearly 70% of striatal PV+ interneurons. Since these interneurons also receive excitatory inputs from the thalamus (Johansson and Silberberg, 2020; Rudkin and Sadikot, 1999; Sciamanna et al., 2015), it is conceivable that glutamate release from thalamic synapses might be sufficient to promote the survival of the remaining striatal PV+ interneurons in our experiments. In contrast, PV+ interneurons are not influenced by the local activity of their main target, the neighboring MSNs, presumably because they receive relatively few inputs from these cells (Chuhma et al., 2011; Taverna et al., 2007).

### Local control of striatal ChAT+ interneuron survival

The final number of striatal ChAT+ interneurons is primarily regulated by the activity of the local MSNs. Increasing MSN activity negatively impacts ChAT+ interneuron survival. This result is consistent with the observation that ChAT+ interneurons receive prominent inputs from these cells (Chuhma et al., 2011; Gonzales et al., 2013; Guo et al., 2015). The neocortex can influence the survival of striatal ChAT+ cells, although this effect is likely mediated by the indirect modulation of MSN activity. Several lines of evidence support this conclusion. First, pyramidal cells provide only weak and sparse direct excitatory inputs to ChAT+ interneurons (Johansson and Silberberg, 2020; Lapper and Bolam, 1992). Second, in the absence of excitatory synaptic drive, MSNs remain quiescent due to a very hyperpolarized resting membrane potential (Wilson and Kawaguchi, 1996). Consequently, in experiments where cortico-striatal inputs are reduced, it is expected that MSN activity would decrease, which would in turn increase the survival of ChAT+ cells. Third, since the number of striatal PV+ interneurons scale up when the number or activity of pyramidal neurons are experimentally raised, increasing cortico-striatal inputs may paradoxically decrease the activity of MSNs, and indirectly increase the survival of ChAT+ interneurons.

It is tempting to speculate on the source of the signals promoting the survival of ChAT+ interneurons. One possibility is that glutamate release from thalamic synapses might contribute towards the survival of these interneurons, as these cells receive more substantial inputs from the thalamus compared to the relatively weak cortical innervation (Johansson and Silberberg, 2020; Lapper and Bolam, 1992). Alternatively, it has been shown that ChAT+ interneurons are tonically active (Aosaki et al., 1995; Wilson et al., 1990). This spontaneous activity is generated by endogenous ionic conductances (Bennett et al., 2000; Wilson, 2005), independently of excitatory synapses (Bennett et al., 2000; Bennett and Wilson, 1999). It is therefore conceivable that the endogenous firing of striatal ChAT+ interneurons might promote the survival of these cells during the period of programmed cell death.

### Outlook

Striatal interneuron dysfunction is associated with neurodevelopmental disorders that affect movement, cognition, and behavior. Alterations in striatal interneuron numbers have been described in patients with Tourette syndrome and schizophrenia (Holt et al., 2005; Kalanithi et al., 2005; Kataoka et al., 2010). Equally, experimental manipulation of the activity and number of these cells leads to motor stereotypies and other functional deficits in mice (Aliane et al., 2011; Crittenden et al., 2017; Rapanelli et al., 2017). In the striatum, the regulation of PV+ interneuron numbers is important for establishing balanced ratios of excitation and inhibition. Furthermore, this balance appears to be crucial for the survival of ChAT+ interneurons. Elucidating whether similar mechanisms operate in humans may shed new light on the neurobiology of neurodevelopmental disorders with alterations in the number of interneurons.

## ACKNOWLEDGMENTS

We thank I. Andrews for excellent technical assistance; S. Anderson (University of Pennsylvania, USA), K. Nave (Max Planck Institute for Experimental Medicine, Germany) and M. Verhage (Vrije Universiteit Amsterdam, The Netherlands) for *Nkx2-1-Cre, Nex^Cre/+^* and *Stxbp1^fl/fl^* mice, respectively; N. Dehorter and B. Rico for critical reading of the manuscript, and members of the Marín and Rico laboratories for stimulating discussions and ideas. Supported by a grant from the European Research Council (ERC-2017-AdG 787355) to O.M.

## Data Availability

All materials, code, and data will be shared upon reasonable request.

## Author Contributions

V.S., K.B. and O.M. conceived the study. K.B. and V.S. performed initial experiments establishing the programmed cell death of striatal interneurons. V.S. performed all experiments described in this manuscript. D.E-A., F.O., C.B., and A.H-G. contributed to cloning and validation of the Cre-out shRNA constructs. E.S-P. and S-E.B. contributed to data collection. R.E. contributed *Vglut1* and *Vglut2* mice. V.S. and O.M. wrote the paper with input from all authors.

## Supplementary Information

**Fig. S1.**
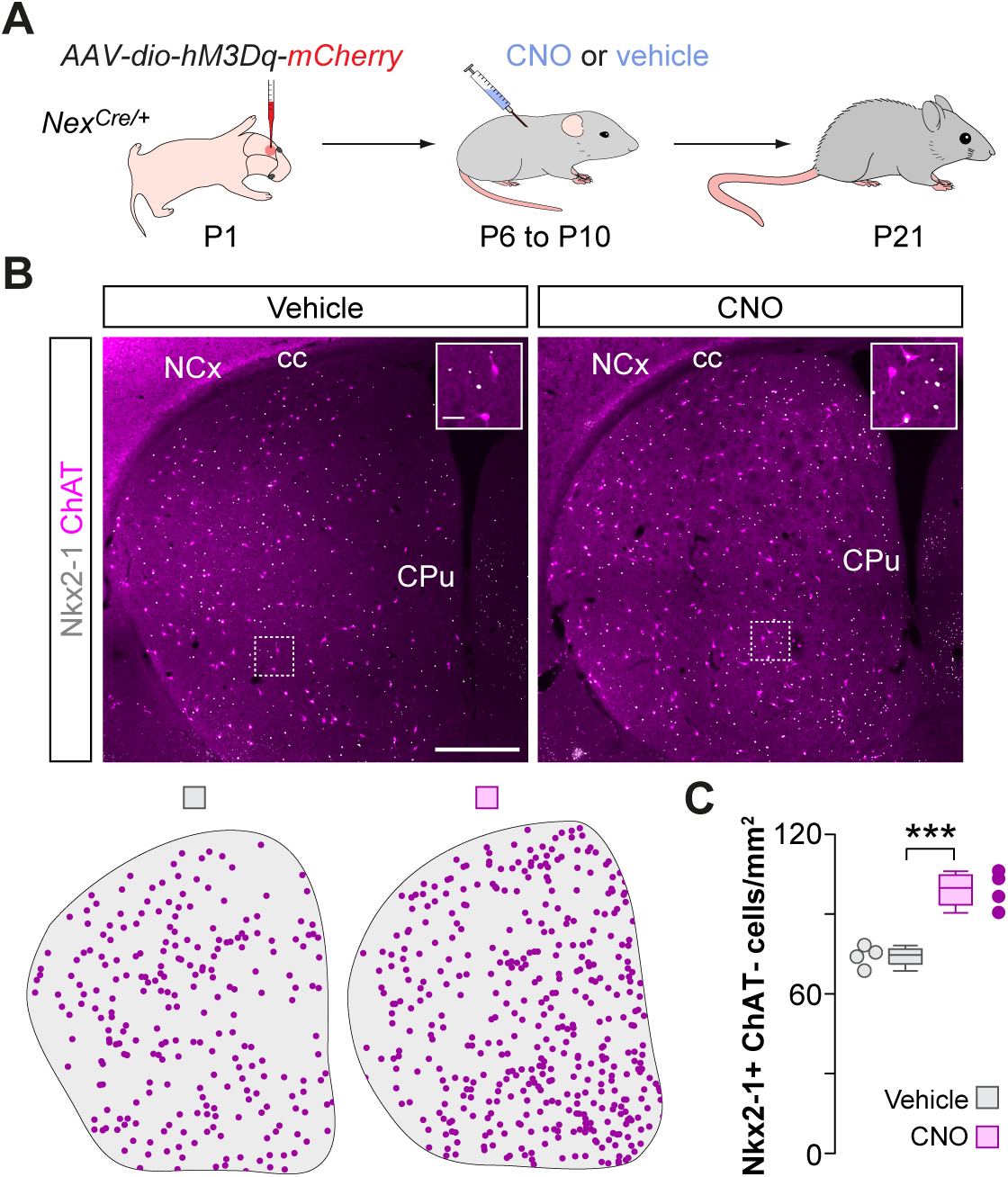
Increase in PV+ interneurons is due to rescue from cell death and not due to activity-related changes in PV+ expression. **(A)** Schematic of experimental design. **(B)** Coronal sections through the striatum of vehicle and CNO treated *Nex^Cre/+^* mice immunostained for Nkx2-1 and ChAT. The schematic dot maps indicate the locations of Nkx2-1+/ChAT-interneurons in each case. **(C)** Quantification of Nkx2-1+/ChAT-interneuron (putative PV+ cell) density in the striatum of vehicle (*n* = 4) and CNO treated (*n* = 4) *Nex^Cre/+^* mice. Two-tailed unpaired Student’s *t*-test, \*\*\**p* < 0.001. Data are shown as boxplots and adjacent data points indicate the average cell density in each animal. Scale bars, 500 *μ*m and 50 *μ*m (inset).

**Fig. S2.**
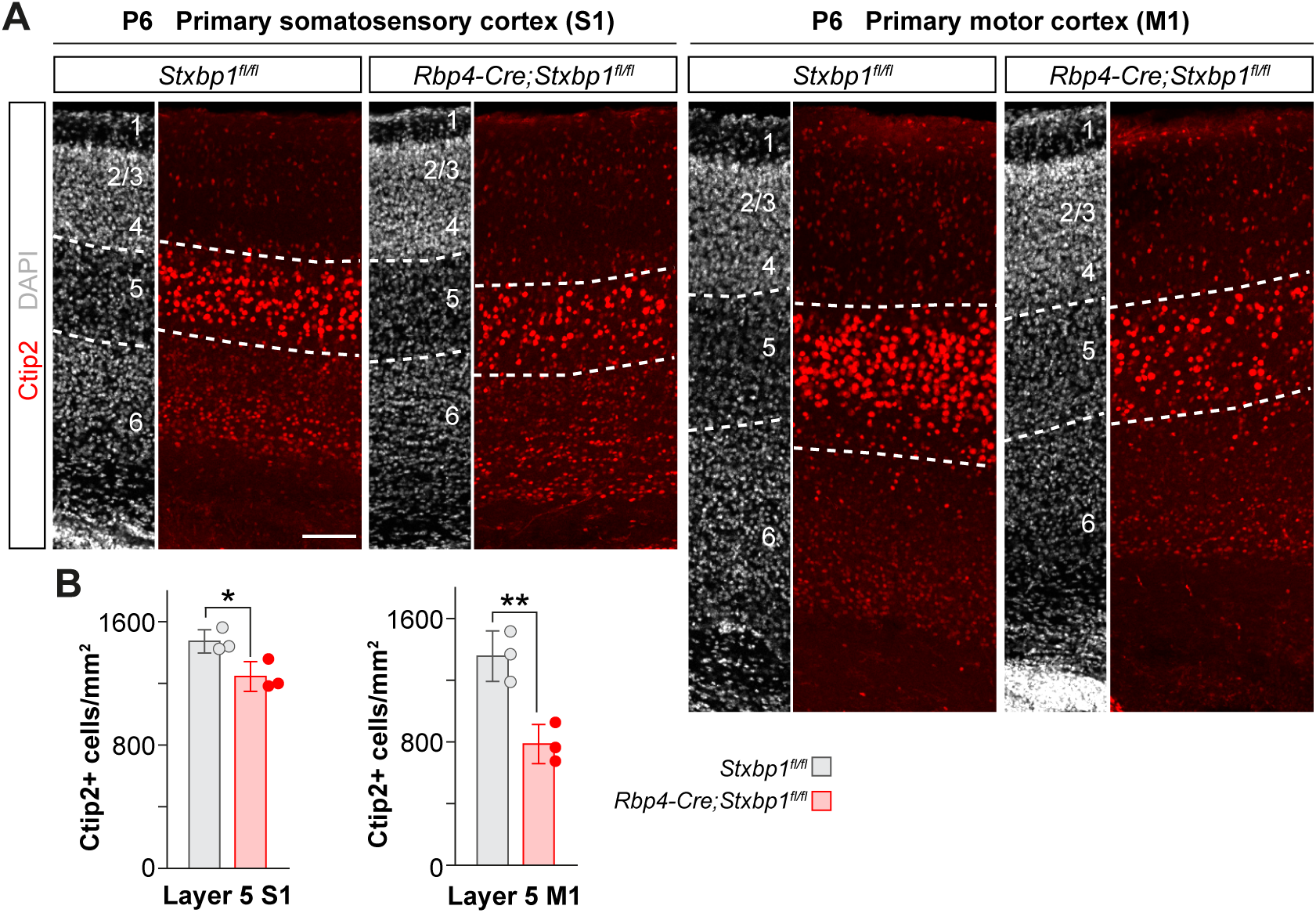
Deleting *Stxbp1* causes the death of layer 5 neurons by postnatal day 6. **(A)** Coronal sections through primary somatosensory cortex (S1) and primary motor cortex (M1) of control *Stxbp1^fl/fl^* and *Rbp4-Cre;Stxbp1^fl/fl^* mice immunostained for Ctip2 at P6. **(B)** Quantification of layer 5 Ctip2+ cell density in S1 and M1 of control *Stxbp1^fl/fl^* (*n* = 3) and mutant *Rbp4-Cre;Stxbp1^fl/fl^* (*n* = 3) mice. S1: two-tailed unpaired Student’s *t*-test, \**p* < 0.05. M1: two-tailed unpaired Student’s *t*-test, \*\**p* < 0.01. Data are shown as bar plots and adjacent data points indicate the average cell density in each animal. Error bars indicate standard deviation. Scale bar, 100 *μ*m.

**Fig. S3.**
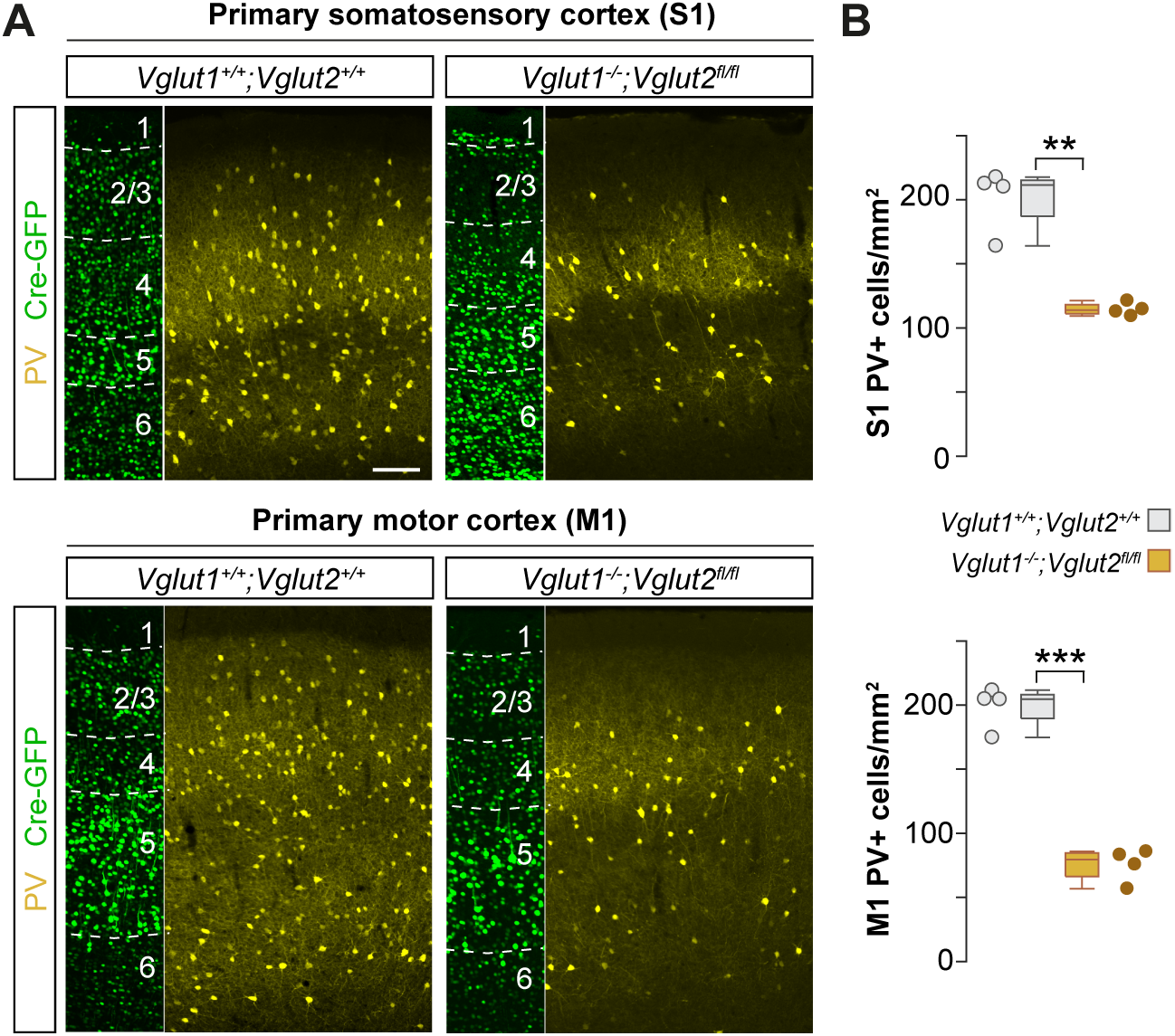
Abolishing cortical glutamatergic transmission impacts the survival of cortical PV+ interneurons. **(A)** Coronal sections through primary somatosensory cortex (S1) and primary motor cortex (M1) of Cre injected control *Vglut1^+/+^;Vglut2^+/+^* and double-mutant *Vglut1^-/-^;Vglut2^fl/fl^* mice. **(B)** Quantification of PV+ interneuron density in S1 and M1 of control *Vglut1^+/+^;Vglut2^+/+^* (*n* = 4) and double-mutant *Vglut1^-/-^;Vglut2^fl/fl^* (*n* = 4) mice. S1: two-tailed unpaired Student’s *t*-test, \*\**p* < 0.01; M1: two-tailed unpaired Student’s *t*-test, \*\*\**p* < 0.001. Data are shown as boxplots and adjacent data points indicate the average cell density in each animal. Scale bar, 100 *μ*m.

**Fig. S4.**
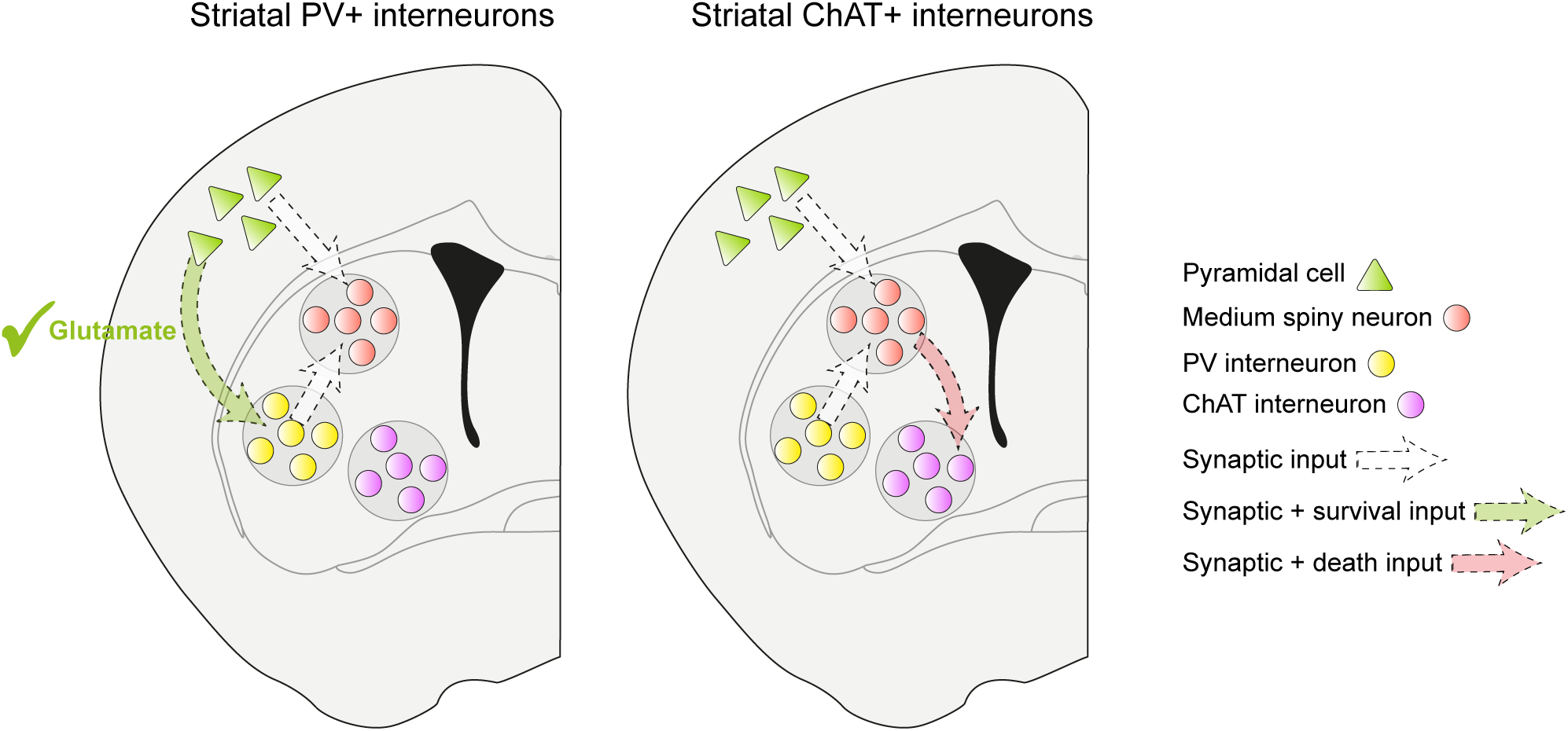
Circuit mechanisms regulating programmed cell death of striatal interneurons. Striatal PV+ interneurons receive glutamatergic input from cortical pyramidal neurons that rescue them from programmed cell death and help to establish appropriate levels of inhibition in the striatum (left). The activity of striatal medium spiny neurons (MSNs) is tightly regulated by a combination of excitatory glutamatergic inputs from the cortex and inhibitory GABAergic inputs from PV+ interneurons. An imbalance leading to hyperactivity in MSNs kills ChAT+ interneurons (right).

## Materials and Methods

### Mice

The mouse lines *Nex^Cre/+^* (*Neurod6^tm1(cre)Kan^*) (Goebbels et al., 2006), *Nkx2-1-Cre* (*Tg*(*Nkx2-1-cre)2Sand*) (Xu et al., 2008), *RCL^tdT^* (*Gt*(*ROSA*)26Sor^tm9(CAG-tdTomato)Hze^) (Madisen et al., 2010), *Bak^-/-^,Bax^fl/fl^* (*Bak1^tm1Thsn^* and *Bax^tm2Sjk^)* (Takeuchi et al., 2005), *Rbp4-Cre* (*Tg(Rbp4-cre)KL100Gsat*) (Gerfen et al., 2013), *Stxbp1^fl/fl^* (*Stxbp1^tm1MVer^*) (Heeroma et al., 2004), *Vglut1^-/-^;Vglut2^fl/fl^* (*Slc17a7^tm1Edw^* and *Slc17a6^tm1.1Thna^*) (Fremeau et al., 2004; Hnasko et al., 2010) were used in this study. Animals were housed in groups of up to five littermates and maintained under standard, temperature-controlled, laboratory conditions. Mice were kept on a 12:12 light/dark cycle and received water and food ad libitum. All animal procedures were approved by the ethical committee (King’s College London) and conducted following European regulations, and Home Office personal and project licenses under the UK Animals (Scientific Procedures) 1986 Act.

### Generation of DNA constructs

The Addgene plasmid *pAAV hSyn-hM3D(Gq)-mCherry* (Addgene 50474) (Roth, 2016) was used as a starting point to generate *pAAV hSyn-lox-hM3D(Gq)-mCherry-lox*. Addgene_50474 was first digested with Sal1 and EcoR1 to generate a 4545 bp backbone for further cloning. The following primers were used to PCR amplify a fragment of 2597 bp containing *hM3D(Gq)-mCherry* flanked by *loxP* sites in the same directional orientation: Fw primer: 5’-CTAGAGTCGACATAACTTCGTATAGCATACATTATACGAAGTTATGCCACCATGACCTTGCAC-3’. Rv primer: 5’-TATCGAATTCATAACTTCGTATAATGTATGCTATACGAAGTTATTTACTTGTACAGCTCGTCC-3’. The fragments were purified using QIAquick gel extraction kit (QIAGEN, Cat# 28704) The PCR amplified fragment was digested with Sal1 and EcoR1 and ligated with the backbone to give the final construct.

### AAV production

AAVs were produced using polyethylenimine (PEI) transfection of HEK293FT cells as described before (Favuzzi et al., 2019). In brief, DNA and PEI were mixed in the ratio of 1:4 in uncomplemented DMEM and left at room temp for 25 mins to form the DNA-PEI complex. The transfection solution was added to each plate and incubated for 72 h at 37 °C in 5% CO2. The transfected cells were then scraped off the plates and pelleted. The cell pellet was lysed in buffer containing 50mN Tris-Cl, 150mM NaCl and 2mM MgCl2 and 0.5% sodium deoxycholate and incubated with 100U/ml Benzonase (Sigma-Aldrich, Cat# E1014 25KU) for 1 h to dissociate particles from membranes. Any remaining contaminants and empty or incomplete viral particles were removed with a discontinuous iodixanol (OptiPrepTM) (Sigma-Aldrich, Cat# D1556) gradient ultracentrifugation using four layers of different iodixanol concentrations of 15, 25, 40 and 58% (Zolotukhin et al., 1999). The viral suspension was loaded on the iodixanol gradient in Quick-seal polyallomer tubes (Beckman-Coulter, Cat# 342414) and spun in a VTi-50 rotor at 50,000 rpm for 75 mins at 12 °C in an Optima L-100 XP Beckman Coulter ultracentrifuge. The recovered virus fraction was purified by first passing through a 100-kDa molecular weight cut off (MWCO) centrifugal filter (Sartorius Vivaspin, Cat# VS2041) and then through an Amicon Ultra 2ml Centrifugal filters (Merck Millipore, Cat# UFC210024). Storage buffer (350 mM NaCl and 5% Sorbitol in PBS) was added to the purified virus and stored in 5 *μ*l aliquots at −80 °C. AAV titer was determined by quantitative polymerase chain reaction (qPCR) using primers for the WPRE sequence that is common to the construct. The following primers were used: WPRE Forward primer: 5’GGCACTGACAATTCCGTGGT-3’. WPRE Reverse primer: 5’-CGCTGGATTGAGGGCCGAAG-3’. The extracted viral DNA and a serial dilution of the transfer plasmid DNA containing the WPRE sequence were transferred to a 96 well plate and measured using a LightCycler^®^ 96 instrument (Roche Life Science). AAVs produced had a titer of 2.10 × 10^13^ viral genomes/ml.

### Stereology

The total number of striatal MGE interneurons was estimated using the optical dissector method (West and Gundersen, 1990)

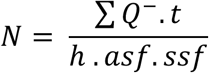

where ΣQ^-^ is the total number of cells counted, *t* the mean section thickness, *h* the height of the optical dissector (18 *μ*m) adjusting for the guard zones (2 *μ*m) above and below the dissector, *asf* stands for the area sampling fraction and *ssf* stands for the section sampling fraction (sampling frequency). An ApoTome (Zeiss) microscope equipped with a motorized stage and color camera was connected to a computer with the Stereo Investigator software (MBF Biosciences). The boundaries of the striatum were first defined with a 2.5x objective (Zeiss) and the entire area was split into 4 equal quadrants.

### Intracranial injections

P1 mice were anesthetized with 3% isofluorane and mounted on a stereotaxic frame. Isoflurane concentration was maintained between 1 and 2% throughout the procedure. Injections of *hSyn-DIO-hM3D(Gq)-mCherry* were targeted to the frontal cortex, at ~ 0.5 mm anterior and 1 mm lateral to bregma, of the left hemisphere. 600 nl of virus were injected at a depth of 400 *μ*m in the left hemisphere at a speed of 3 nl sec^-1^. Injections of *hSyn-DIO-hM4D(Gi)-mCherry* were targeted to 6 sites in the left hemisphere to infect layer 5 neurons in the entire cortex. At each site, 200 nl of virus were injected at a depth of 600 *μ*m at a speed of 3 nl sec^-1^. Injections of *hSyn-GFP-Cre* were targeted to 6 sites in the left hemisphere to infect neurons in the entire cortex. At each site, 200 nl of virus were injected at a depth of 400 *μ*m at a speed of 3 nl sec^-1^. Injections of *hSyn-lox-hM3D(Gq)-mCherry-lox* were targeted to a single site in the left striatum. Along the anteroposterior axis, the site was located halfway between bregma and the blood vessel lying immediately posterior to the olfactory bulb and at ~1.3 to 1.5 mm lateral to the midline. 600 nl of virus were injected at a depth of ~1.4 to 1.6 mm at a speed of 3 nl sec^-1^ to infect neurons in the dorsal striatum.

### CNO injections

Injected animals were randomly assigned to the “Vehicle” or “CNO” groups. Clozapine-N-oxide (Tocris Bioscience, Cat# 4936) was dissolved in 0.9% saline and 0.5% DMSO (Sigma-Aldrich, Cat# D8418) to give a final concentration of 0.5 mg ml^-1^. The vehicle consisted of 0.5% DMSO in saline without the active ligand. In experiments where the activating DREADD hM3D(Gq) was expressed in cortex, CNO or Vehicle was administered subcutaneously twice a day from P6 until P10 at a dose of 1 mg kg^-1^. This period was chosen since P6 and P10 correspond to the start and end of striatal interneuron death, respectively. In experiments where the hM3D(Gq) was expressed in the striatum, CNO or Vehicle was administered subcutaneously thrice a day from P6 until P10 at a dose of 5 mg kg^-1^. In experiments where the inactivating DREADD hM4D(Gi) was expressed in cortex, CNO or Vehicle was administered subcutaneously thrice a day from P6 until P10 at a dose of 5 mg kg^-1^.

### Histology and immunohistochemistry

Mice were deeply anesthetized with pentobarbital sodium (Euthatal, Merial Animal Health Ltd) by intraperitoneal injection, and were transcardially perfused with 0.9% NaCl solution (Sigma-Aldrich, Cat# S76530) followed by 4% paraformaldehyde (PFA) (Sigma-Aldrich, Cat# 441244) dissolved in phosphatase-buffered saline (PBS). Brains were extracted and post-fixed for 2h at 4°C. The brains were immersed overnight in 30% sucrose (Sigma-Aldrich, Cat# S0389) and coronal sections were cut frozen on a sliding microtome (Leica SM2010 R) to a thickness of 40 μm. Every 8th section was used for antibody staining. The sampling was therefore every 280 *μ*m. Free-floating brain slices were permeabilized with 0.25% Triton X-100 (Sigma-Aldrich, Cat# T8787) in PBS for 30 mins, and blocked for 1h in a blocking buffer containing 0.25% Triton X-100, 10% normal donkey serum (Sigma-Aldrich, Cat# S30), and 5% bovine serum albumin (BSA) (Sigma-Aldrich, Cat# A8806). Primary antibodies were dissolved in fresh blocking buffer and the sections were incubated overnight at 4°C in the primary antibody solution. The following day, the tissue was repeatedly rinsed. Secondary antibodies were dissolved in fresh blocking buffer and the sections were incubated in the secondary antibody solution for 2h at room temperature. The sections were counterstained with 5 *μ*M 4’,6-diamidine-2’-phenylindole dihydrochloride (DAPI) (Sigma-Aldrich, Cat# D9542) in PBS, rinsed repeatedly and mounted on a glass slide with Mowiol and DABCO. The following primary antibodies were used: Rabbit anti-parvalbumin (1:1000, Swant, Cat# PV-27), goat anti-ChAT (1:500, Millipore, Cat# AB144P), rabbit anti-TTF1 (1:500, Abcam, Cat# ab76013), rat anti-Ctip2 (1:1000, Abcam, Cat# ab18465). The following secondary antibodies were used: Donkey-anti-rabbit-Alexa 555 (1:400, Molecular Probes, Cat# 31572), donkey anti-rabbit-Alexa 488 (1:400, Thermo Fisher Scientific, Cat# 21206), donkey anti-rabbit-Alexa 647 (1:400, Thermo Fisher Scientific, Cat# 31573), donkey-anti-goat Alexa 488 (1:400, Molecular Probes, Cat# 11055), donkey-anti-goat Alexa 555 (1:400, Invitrogen, Cat# 21432), donkey-anti-goat Alexa 647 (1:400, Thermo Fisher Scientific, Cat# 21447), donkey-anti-rat Cy3 (1:400, Jackson ImmunoResearch Europe Ltd., Cat# 712-165-150).

### Image acquisition and image analysis

Tile scan images of the striatum were acquired at 1,024×1,024-pixel resolution and 8-bit depth on an inverted Leica TCS-SP8 confocal microscope. 7 to 8 sections covering the striatum along the rostrocaudal axis were imaged. Samples from the same experimental litter were imaged and analyzed in parallel, using similar laser powers, photomultiplier gain and detection filter settings. Images were analyzed using custom routines written in MATLAB. Briefly, images were first adjusted for brightness and contrast. A median filter was applied to the image to remove salt-and-pepper noise followed by Sobel filtering to determine the edges of the cells. This was followed by the following morphological operations: one round of dilation (*imdilate* function), one round of filling (*imfill* function) and two rounds of erosion (*imerosion* function). The cell positions were calculated as the centroids (*regionprops* function). The striatum was delineated and the number of centroids inside it was counted and divided by the total area of the region to give the cell density in cells per mm^2^. This analysis was repeated for all sections where the striatum was present and individual values for the sections were averaged to give a single value per animal.

### Statistical analysis

All statistical analyses were performed using MATLAB (Supplementary Table 1). Group data are presented as boxplots. On each box, the central mark indicates the median and the top and bottom edges of the box indicate the 25th and 75th percentiles respectively. The whiskers extend to the data points not considered as outliers. Outliers were considered as points lying at 1.5 x IQR (inter-quartile range) above the 75th percentile or under the 25th percentile, thereby giving coverage of ~99%. The Anderson-Darling normality test was used to compare the empirical distribution of the data sets with a normal probability distribution. To compare normally distributed data, we used paired or unpaired two-tailed Student’s t-test. To compare non-normally distributed data, we used Wilcoxon’s rank-sum test for unpaired comparisons or Wilcoxon’s signed-rank test for paired comparisons respectively. To analyze the differences among multiple experimental groups, we used a one-way analysis of variance (ANOVA) followed by the Tukey-Kramer post-hoc test. Statistical significance was considered at p-values ≤ 0.05. The number of animals for each experiment is described in each figure legend.

